# Mate choice and gene expression signatures associated with nutritional adaptation in the medfly (*Ceratitis capitata*)

**DOI:** 10.1101/362210

**Authors:** Will Nash, Irina Mohorianu, Tracey Chapman

## Abstract

Evolutionary responses to nutrition are key to understanding host shifts and the potential for reproductive isolation. Previously, experimental evolution was used to describe responses of the medfly (*Ceratitis capitata*) to divergent larval diets. Within 30 generations there was divergence in larval development time, egg to adult survival and adaptation in adult body size. In this study, the gene expression signatures associated with these changes were examined, using mRNA-seq on males following 60 generations of nutritional selection. Variation in gene expression was also validated using qRT-PCR. The results showed observed an over-representation of differential expression in metabolism, oxidative phosphorylation and proteolysis genes. In addition, at generations 60 and 90, we tested whether these evolved patterns (differences in gene expression) were associated with mate choice. We found evidence for assortative mating by diet at generation 60, but not in mating tests within and between replicate lines at generation 90. Hence, there was no consistent evidence for assortative mating by diet, which emphasises the importance of replicate tests of evolutionary responses over time. Overall, the study provides the first genome-wide survey of the putative mechanisms underpinning evolved responses to nutritional adaptation.

## Introduction

Adaptation to different nutritional ecologies can play key roles in successful expansion into new habitats as well as in reproductive divergence associated with host shifts with the potential to result in reproductive isolation (RI) (Schluter 2000; Coyne and Orr 2004; Rundle and Nosil 2005). For example, in *Rhagoletis* fruit flies (Feder et al. 1994) genetic divergence between populations is associated with host plant shifts. If the host shift occurs in sympatry, this can lead to the subsequent formation of host races (Linn et al. 2003; Nosil et al. 2012; Powell et al. 2014; Soria-Carrasco et al. 2014). This local adaptation and host race formation can have a significant impact on traits associated with fitness on the new host and also, potentially, RI (Feder et al. 1994; Dambroski and Feder 2007; Nosil 2007; Nosil et al. 2012). However, much is still unknown concerning the instigation, and underlying mechanisms, of nutritional adaptation (Panhuis et al. 2001; Maan and Seehausen 2011; Butlin et al. 2012; Safran et al. 2013).

Mechanistic insights into nutritional adaptation can be gained from next generation sequencing (NGS), which is revolutionizing the study of natural selection (Seehausen et al. 2014), adaptation (Elmer and Meyer 2011; Savolainen et al. 2013) and the initial steps of speciation (Nosil 2012). The characterization of patterns of gene expression across genomes offers new opportunities for the study of the putative causal links between the differences in gene expression and adaptive genetic divergence (Pavey et al. 2010). For example, populations of *Drosophila mojavensis* from different regions of the Sonoran desert exhibit premating isolation, mediated by cuticular hydrocarbon profiles (CHCs) and courtship song, based on the cactus host they utilise for reproduction (Etges et al. 2007, 2009; Etges 2014). Transcriptomic studies have identified a small suite of metabolic, olfactory and behavioural trait genes whose expression is directly influenced by the host food plant (Matzkin 2012; Rajpurohit et al. 2013; Etges et al. 2015). NGS methods can also be combined with experimental evolution for additional insights into divergence (Rice and Hostert 1993; Kawecki et al. 2012) particularly via ‘*evolve & resequence’* techniques (Schlötterer et al. 2015). By imposing selection on a particular trait or suites of traits, experimental evolution studies can reveal adaptive responses to selection in real time (Remolina et al. 2012).

Here we used mRNA-seq to capture the effects of selection mediated by dietary adaptation on protein-coding gene expression in an experimental evolution framework. Experimental evolution was conducted in the medfly (*Ceratitis capitata*), a global pest of significant agricultural importance, under two divergent larval dietary treatments, to characterise the responses to nutritional selection (Leftwich et al. 2017) and to describe the underlying changes in expression associated with these responses.

The medfly is an extreme generalist and exhibits wide plasticity in host selection, utilization (Levinson et al. 1990; Yuval and Hendrichs 2000) and oviposition (Prokopy et al. 1984; Yuval and Hendrichs 2000). Larvae are viable in a wide range of fruits and diets, from both inside and outside of the known host range (Carey 1984; Krainacker et al. 1987; Zucoloto 1993; Nash and Chapman 2014). This highly plastic host choice behaviour is evident in reports that the medfly can infest over 350 different host fruits (Liquido et al. 1991). As such, the medfly is of huge economic importance as an agricultural pest. Its ability to exhibit strong plasticity (Carey 1984; Nyamukondiwa et al. 2010) is thought to have facilitated its global invasion (Gasperi et al. 2002; Malacrida et al. 2007). Global populations of medfly vary in a wide range of demographic (Diamantidis et al. 2008a,b, 2009, 2011a) and behavioural traits (Briceño et al. 2002, 2007), as well as in genetic structure (Gasperi et al. 2002; Karsten et al. 2015). However, unlike for some Drosophilid species (e.g. Dodd 1989; Rundle et al. 2005), there are as yet no reports of mating isolation between different strains exhibiting different diets (Cayol et al. 2002). In general, mechanisms influencing nutritional plasticity are recognised as key to understanding nutritional adaptation. However, as yet we know little about the underpinning genetic basis and regulatory interactions. This is the key gap investigated in this study.

Experimental evolution was practised on two sets of three independent evolutionary replicates, each selected on different larval diets with different nutritional complexity and caloric value (Leftwich et al. 2017). Within 30 generations, this dietary selection led to divergence in larval development time and egg to adult survival and the emergence of local adaptation in adult body size (Leftwich et al. 2017). In the current study, we investigated the evolved gene expression signatures associated with this divergence following 60 generations of selection. We used mRNA-seq to compare gene expression between sexually mature males, following rearing for 2 generations on a common garden diet. To capture tissue specificity, we profiled the Head/Thorax and Abdomen separately. Evolved gene expression patterns were highly replicable, 214 transcripts were identified as differentially expressed (DE) between the dietary treatments. DE was observed in a suite of genes involved in nutrient metabolism, oxidative phosphorylation (OXPHOS), and proteolysis. Nutritional adaptation was associated with replicable but inconsistent differences in mating preferences at generations 60 and 90, suggesting no underlying assortative mating by diet as a by-product of adaptation to divergent selection on larval diets.

## Materials and Methods

### EXPERIMENTAL EVOLUTION LINES

The experimental evolution lines were derived from the TOLIMAN wild type (originating from Guatemala and reared in the laboratory since 1990; Morrison et al. 2009). For at least two years prior to the start of these experiments the TOLIMAN strain was reared on a wheat bran diet (24% wheat bran, 16% sugar, 8% yeast, 0.6% citric acid, 0.5% sodium benzoate). To initiate the experimental evolution, flies were established on (i) sucrose-based ‘ASG’ medium (1% agar, 7.4% sugar, 6.7% maize, 4.75% yeast, 2.5% Nipagin (10% in ethanol), 0.2% propionic acid, 684kcal/L) or (ii) ‘Starch’ (S) medium (1.5% agar, 3% starch, 5% yeast, 0.4% propionic acid, 291 kcal/L) (Leftwich et al. 2017). Three independent biological replicates of each of the two regimes were maintained under allopatry. All experiments and culturing was conducted at 25°C, 50% relative humidity, on a 12:12 light dark photoperiod. Adults emerging from each replicate were maintained in groups of approximately 30 males and 30 females in plastic cages (9cm x 9cm x 9cm). Adults from all lines received the same standard adult diet (*ad libitum* access to sucrose-yeast food; 3:1 w/w yeast hydrolysate: sugar paste and water). Each generation, approximately 500 eggs were placed on 100ml of the appropriate diet in a glass bottle. When third instar larvae started to ‘jump’ from the larval medium, the bottles were laid on sand and pupae allowed to emerge for seven days. Pupae were then sieved from the sand and held in 9mm petri dishes until eclosion of adults began and the next generation was initiated.

### IDENTIFICATION OF GENE EXPRESSION SIGNATURES OF NUTRITIONAL ADAPTATION USING MRNA-SEQ

Eggs were derived from the ASG (A) and Starch (S) experimental evolution lines at generation 60 and reared on a common garden glucose (CG) diet (1.5% agar, 3% glucose, 5% yeast, 0.5% propionic acid) under standard conditions and density for two generations. Adults emerging from daily collected cohorts of pupae were sex-sorted and reared in single sex cages until seven days post eclosion. 17 - 30 male flies were then flash frozen in liquid N_2_ 30 minutes after lights on (09:30), in Eppendorf tubes, in groups of 10 - 15. These samples were then stored at −80°C until RNA extraction.

### RNA EXTRACTION AND MRNA-SEQ

Prior to RNA extraction, males were split into Head/Thorax (HT), and Abdomen (Ab), over dry ice. Total RNA was extracted from samples of 17 - 22 body parts pooled within each replicate of each dietary treatment using the *mir*Vana kit (Ambion), according to the manufacturer’s instructions. 5µg (>200ng/µl) of Total RNA from male flies was submitted for sequencing. mRNA-seq was conducted (BaseClear provider) on libraries prepared using the Illumina TruSeq protocol, following polyA selection for mRNA. Sequencing was single-end, 50 cycles on the Illumina HiSeq 2500 platform (Rapid Mode).

### BIOINFORMATICS

#### Quality control

Initial quality control was performed on original FASTQ files. FASTQ files were then converted into FASTA format. Reads containing Ns (< 1%) were discarded. The set of redundant reads was collapsed into non-redundant format and the complexity (ratio of non-redundant reads to redundant reads) calculated (Mohorianu et al. 2011, 2017a). To evaluate the extent of the sequencing bias, the per base nucleotide composition was determined. The proportion of reads incident to the reference genome and available annotations was calculated before and after normalization of expression levels, to ensure that the normalization targeted technical inter-sample variability without altering the biological features present (Mohorianu et al. 2017a).

#### Read mapping and normalisation

The mapping of all reads was done using PaTMaN (Prüfer et al. 2008), full length, with no gaps allowed and with maximum 1 mis-match at any position along the read. For this analysis, we used the Ccap1.1 version of the *Ceratitis capitata* genome and curated *C. capitata* transcripts downloaded from NCBI (for rRNAs and protein coding genes). rRNA-reads were excluded from all samples and we then sampled the filtered reads to a fixed total of 29M (the minimum sequencing depth, post filtering), using subsampling (without replacement) (Mohorianu et al. 2017a). Gene expression was calculated as the algebraic sum of the abundances of incident reads. Any remaining variability in the expression matrix was then minimised by using quantile normalisation (Bolstad et al. 2003). The proportion of reads incident to the mRNA annotations corresponding to v1.1 of the *C. capitata* genome was significantly lower than the proportion of genome matching reads. Hence, we used curated *C. capitata* transcripts downloaded from NCBI, which yielded comparable proportion of transcript- and genome-matching reads.

#### Differential expression analysis

We first investigated differential expression (DE) between replicates, using a log_2_ offset fold change (OFC, with empirically-determined offset of 20 (Mohorianu et al. 2017a) to filter out low level noise. Transcripts with a normalized abundance of <100 were excluded. The DE analysis was using a hierarchical approach on two levels (Mohorianu et al. 2017b): (i) body part (tissue) (HT/Ab) and (ii) dietary treatment (A/S). The DE call between treatment types was made on maximal confidence intervals (CIs) and genes with a log_2_(OFC) > 1 (corresponding to a 2-fold change difference) were called DE.

#### Annotation and functional description

To expand the annotation information, DE transcripts were matched to their corresponding *Drosophila melanogaster* homologues using tBLASTx (Altschul et al. 1990). The similarity threshold for matching was alignment length > 50nt, with similarity > 50%. The similarity threshold following matching was alignment length > 50nt, with similarity > 50%. The tissue-specific gene lists resulting from this were tested for functional enrichment using g:Profiler (Reimand et al. 2016) using all *D. melanogaster* genes as the background set.

### QRT-PCR VALIDATION

Candidates for validation were chosen based on their expression levels (>200 normalized expression) and to cover representative GO categories. Presence plots were used to enhance primer design (i.e. to avoid primers spanning exon/exon boundaries). We first assessed the stability of expression of 3 reference genes (Scolari et al. 2012; Koramutla et al. 2016) in our RNA-seq data. We selected 3 reference genes *(CcRpL13a, CcRpL27*, *poll*) based on a circular permutation test, high read abundance (> 500,000 reads in each sample, in both body parts) and by assessing the consistency of expression between tissues (|DE| <= 0.0001) and treatments. We then designed primers (using primerBLAST (Ye et al. 2012)) and checked, using PatMaN (Prüfer et al. 2008) for potential multiple matches to other genomic locations (primer details in Table S1). We chose 15 candidate genes of interest (GOIs) for validation.

For primer optimisation we used mRNA extracted from whole wild type flies. We removed DNA using TURBO™ DNase (Life Technologies) on 2 μg total RNA. DNase was deactivated using the resin in the TURBO™ DNA-free kit (Life Technologies) and all samples were tested for DNA contamination with no-reverse transcription controls. The QuantiTect Reverse Transcription Kit (Qiagen) was used to reverse transcribe 1 μg total RNA to cDNA, which was then stored at −20 °C. qRT-PCRs were run using a StepOnePlus™ machine (Life Technologies) using iTaq Universal SYBR^®^ Green Supermix with triplicate technical replicates using 10 ng RNA as template in 20 μl reactions in MicroAmp^®^ Fast Optical 96-Well Reaction Plate with Barcode, 0.1 ml (Life Technologies). The qRT-PCR conditions were 95 °C for 30 seconds followed by 40 cycles of 95 °C for 15 seconds and 60 °C for 1 minute and data acquisition. Following the qRT-PCR we ran a melt curve analysis (on default settings). All samples showed a single peak on the melt curve. We included a no template control for each GOI on each plate. All primers were used at a final concentration of 5 μM. For primer optimisation (Eurofins MWG Operon) we produced standard curves using at least five 1:5 dilutions of RNA starting at 50 ng cDNA. Samples where no signal was detected during qRT-PCR were given a nominal CT value of 40, to allow statistical testing. The difference in 2^-ΔCT^ between treatments was tested using a two-sample t-test for each GOI.

### TESTS FOR ASSORTATIVE MATING BY DIET

We conducted mate choice tests at generation 60 and 90. Eggs were seeded onto a common garden glucose diet two generations prior to testing (glucose diet: 1.5% agar, 3% glucose, 5% yeast, 0.5% propionic acid). All mating tests were conducted simultaneously. At generation 90, we conducted the same mate choice tests but in addition, examined tests between replicates within each dietary treatment. Flies were sorted by sex within 24 hours of eclosion to ensure virginity. Experimental flies were reared in standard 0.8L rearing cages. To enable identification, one male and one female from each population in each mating test were marked with a spot of red paint on the dorsal side of the thorax. Treatments were fully controlled for handling effects and paint marking.

#### Assortative mating assays

A multiple-choice design was employed in order to maximise the opportunity for mate choice. Five days post eclosion and 48 hours prior to the mating tests, 25 females of each of the two treatments, and similarly 25 males of each treatment, were placed into two, single sex, 0.8L rearing cages. The two cages were connected via a sliding door. Both cages were supplied with *ad libitum* 3:1 sugar yeast hydrolysate diet and water. Mating tests were conducted when the flies were 7-8 days post eclosion, with sexes and treatments exactly balanced for age composition. Mating tests were initiated at 09:30, 30 minutes after lights on, by slowly raising the sliding door. Mating pairs were gently removed and placed in numbered 1.5ml eppendorf tubes for later identification. Three replicates were conducted on each test day, starting at 30 min intervals. The order in which the replicate pairs were tested was alternated, to control for effects of different start times. Each assay continued until 25 mated pairs had been collected, or until 30 minutes had elapsed. The collection of 25 pairs amounted to half of the total population in the cages. Therefore, any effect of diminishing choice due to removal of flies from the cage was minimized (Casares et al. 1998). At generation 60, four replicates were conducted for each combination of tests. At generation 90, five replicates each of 16-25 staring individuals of each sex per line were tested for the own diet and common garden mating tests. For the within line mating tests, four replicates of 25 starting individuals of each sex were tested for the A v A mating assays, and three replicates of 19-25 individuals of each sex per line were tested for S v S. The identity of both individuals in each mating pair was recorded. As only 50% of matings were sampled, mating pairs were treated as independent (Casares et al. 1998; Coyne et al. 2005) and results were pooled by line replicate prior to further analysis.

### STATISTICAL ANALYSIS

The number of observed and total possible pairings for each pair type was calculated. These raw data were then analysed using JMATING v1.0 (Carvajal-Rodriguez and Rolan-Alvarez 2006) to calculate descriptive coefficients based on the cross product estimator of isolation (Rolán-Alvarez and Caballero 2000). *I*_PSI_, a joint isolation index, varying from −1 to 1 (with +1 being total assortative mating, −1 total dissasortative mating and 0 random mating; Coyne et al. (2005)) was used to describe total isolation. The PTI coefficient described positive and negative preferences for mating pairs within each line pair, at each time point. An index of mating asymmetry (*IA*_PSI_) captured the difference in frequency between homotypic and heterotypic pairs (Rolán-Alvarez 2004). Values of *IA*_PSI_ centre around 1 (no asymmetry), with values below one reflecting asymmetry towards the first pair type, and > 1 representing asymmetry towards the second. Finally, we calculated *W*, the cross product estimator of sexual selection, to compare the sexual fitness of males and females (Carvajal-Rodriguez and Rolan-Alvarez 2006) and to give the relative fitness of each treatment in comparison to the fittest treatment within each line pairing. The significance of PSI, PSS, and PTI coefficients was determined as the bootstrap probability of rejecting the null hypothesis of random distribution, after 10,000 iterations of resampling (Rolán-Alvarez and Caballero 2000). When applied to the isolation index (*I*_PSI_), the asymmetry index (*IA*_PSI_), and the estimator of sexual selection (*W*), this is the two-tail bootstrap probability that the value is significantly different from one (i.e. random mating, or zero asymmetry). To test for the overall effect of diet in RI as indicated by *I*_PSI_, *IA*_PSI_ or *W*, probability values for each line replicate comparison were combined using Fisher’s sum of logs method, implemented with the ‘metap’ package in R (Dewey 2016). All other data handling and statistical analysis was conducted in R (R Development Core Team 2015, ver 3.1.1).

## Results

### MRNA-SEQ

#### Quality control

Raw FASTQ files contained between 34.3M and 53.2M reads. Following conversion to FASTA format, we accepted > 99% reads for every sample (Table S2). Other initial checks, such as the nucleotide composition across reads, revealed the expected RNA-seq bias, which was consistent for all samples (Mohorianu et al. 2017a). Next, to optimize the alignment step, the files were transformed from redundant to non-redundant format, yielding between 7.6M and 10.2M unique (non-redundant) reads, with a complexity of between 0.192 and 0.23. The tight range of complexities indicated a reliable sequencing output. The variation in complexities observed between replicates, derived primarily from variation in sequencing depth, suggested that a subsampling normalization would be appropriate.

#### Genome matching and normalisation of expression levels

First, rRNA mapping reads were excluded. Next, the resulting reads were subsampled at a fixed total (29M). The non-redundant read count varied between 7M and 8.2M reads, with complexity between 0.235 and 0.277, showing that the variation in complexity was reduced following normalization (Table S2). On the normalized data, the proportion of reads matching to the ‘Ccap1.1’ genome was: redundant 80 to 83%, non-redundant 77%. The proportion of redundant reads incident to the genome was in line with the results observed for the original data, suggesting that the normalization successfully minimised technical variability. The proportion of reads matching to Ccap NCBI transcripts was: redundant 75 to 83%, non-redundant 75 to 79% (Table S2). This was similar to the proportion of genome matching reads, a result that was expected as polyA selection was used for the sequencing of the mRNA fragments. As the Ccap NCBI transcript annotations provided a similar proportion of R / NR mapping reads to the reference genome, this dataset was used for the subsequent analysis.

### IDENTIFICATION OF DIFFERENTIALLY EXPRESSED PROTEIN-CODING GENES

The normalisation provided comparable distributions of expression across samples, with good agreement between replicates throughout (Fig. S1). The amplitude and frequency of DE (Table S3A-D) between tissues (black line, Fig. S2) was greater than for diets treatment (red line, Fig. S2) showing the hierarchical order of levels within the distribution of DE. A total of 359 transcripts showed DE between treatments (Table 1), with 194 being expressed higher in Starch flies, and 161 higher in ASG. These transcripts were matched to *D. melanogaster* homologues using tBLASTx, which yielded a total of 94 matching genes, 29 in expressed in the HT tissue and 65 in Ab. Tests for functional enrichment of this set highlighted 3 enriched GO terms and 1 KEGG pathway, associated with 15 *C. capitata* transcripts (Fig. 1).

**Table 1.**
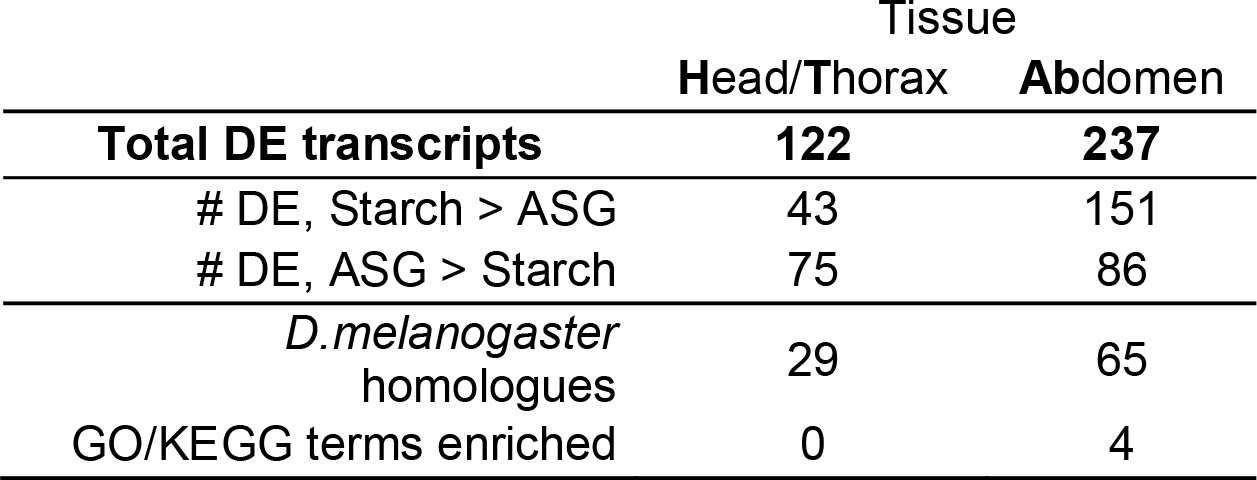
Differentially expressed (DE) transcripts used for functional enrichment. DE between treatment types was called on maximal confidence intervals, with genes exhibiting a log_2_(OFC) > 1 (corresponding to a 2-fold change difference) classed as DE. From this set tBLASTx (Altschul et al. 1990) was used to retrieve homologues from *Drosophila melanogaster*. These homologues were functionally enriched using g:Profiler (Reimand et al. 2016). The number of transcripts which were retained in the analysis at each stage, and the resulting number of enrichment terms are displayed per tissue.

**Figure 1.**
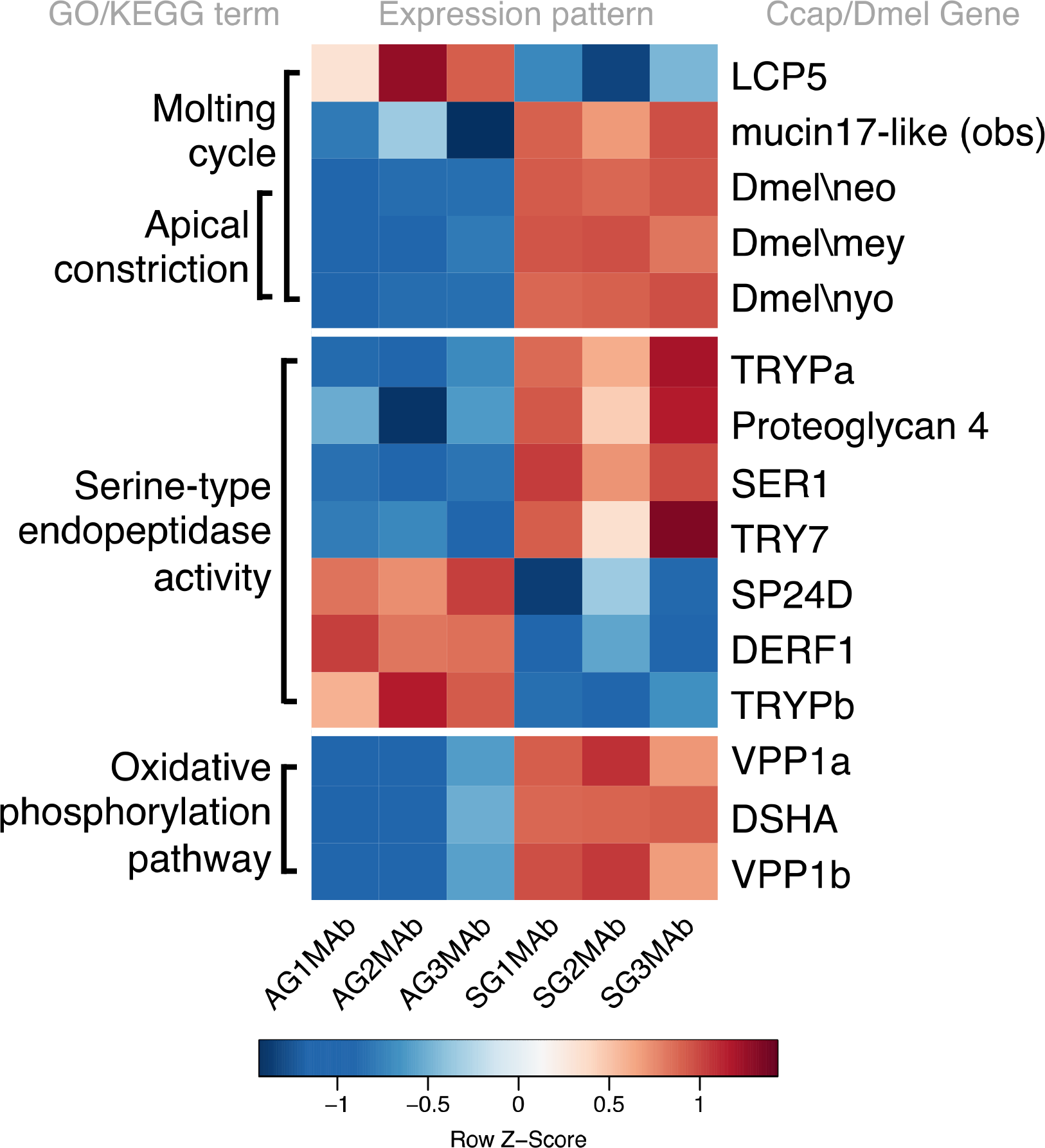
Differentially expressed transcripts in Abdominal tissue of males after 60 generations of adaptation to divergent larval diets. Colour represents row wise Z-score of expression levels normalised by row mean and standard deviation, red represents comparative up regulation, blue represents comparative down regulation. Groupings based on GO enrichment of *D. melanogaster* homologues of medfly DE transcripts are displayed on the left side of the heat map. Gene names are displayed on the right-hand side. ASG samples (AG) are displayed in the 3 left hand columns of the heat map, Starch samples (SG) in the 3 right hand columns, each for replicates 1-3, male abdomen (MAb).

### DIFFERENTIAL EXPRESSION IN THE HEAD+THORAX

Of the 122 transcripts that yielded a signature of DE in the HT, 43 were expressed at higher levels in ASG flies, and 75 higher in Starch (Table S3A, B). 29 of these were matched to *D. melanogaster* identifiers following tBLASTx searching. Functional enrichment did not return any significant terms. After removal of duplicates present due to isoform variation, 65 DE transcripts remained (Table S3B). Of these, 22 bore informative NCBI predicted annotations and 9 were predicted as uncharacterised loci. As examples from this set, homologues for the *D. melanogaster* serine protease genes γTry and CG34458, as well as cytochrome P450 genes Cyp6a9 and Cyp6a21 were expressed at higher levels in Starch flies. Consistent with this, several transcripts predicted to be serine proteases also showed higher expression in Starch flies (e.g. XM_004517825.1, XM_004517721.1). Homologues of the developmentally associated larval cuticle protein genes Lcp3 and Lcp56Ab1 were expressed at higher levels in ASG flies. The functional significance of the enrichment for serine proteases in Starch and cuticle genes in ASG flies is not yet clear.

### DIFFERENTIAL GENE EXPRESSION IN THE ABDOMEN

237 transcripts were DE in abdomen, with 151 expressed higher in Starch flies and 86 higher in ASG (Table S3C, D). Following tBLASTx, 65 *D. melanogaster* identifiers were retrieved. Functional enrichment of these revealed significant overrepresentation for genes involved in the molting cycle (GO-BP:0042303, *P* = 0.0432) and apical constriction (GO:0003383, *P* = 0.0493). Additional enrichment was observed in the molecular function GO term serine-type endopeptidase activity (GO:0004252, *P* = 0.0399) and in genes associated with the oxidative phosphorylation KEGG pathway (ko00190, *P* = 0.0495; Fig. 1). The biological process (BP) ‘molting cycle’ was associated with 5 *C. capitata* genes. Of these, the majority were expressed at a higher level in Starch flies (Fig. 1). However, a single *C. capitata* gene (LCP5) showed the opposite pattern and was expressed at higher levels in ASG flies. This pattern of LCP5 expression was also seen in the HT for ASG flies. Of the five genes enriched for molting cycle, 3 were also enriched for apical constriction. These genes were all expressed at higher levels in Starch than in ASG flies. For all three of these transcripts there were no *C. capitata* annotations, and hence these genes are referred to by their *D. melanogaster* identifiers in Fig. 1. The majority of genes associated with a functional enrichment were serine-like peptidases. 7 *C. capitata* transcripts were associated with this GO term, and exhibited a bidirectional expression pattern, with 4 expressed at higher levels in Starch than in ASG, and the remaining 3 showing the opposite pattern. The final functional enrichment information came from 3 *C. capitata* transcripts that were associated with the KEGG pathway describing oxidative phosphorylation, these were all expressed at higher levels in Starch flies.

After duplicates were removed, 152 abdomen transcripts showing DE remained (Table S3B). Of these, 53 were associated with *Ceratitis capitata* annotations or predicted annotations and 21 were predicted as uncharacterised *C. capitata* loci. Within this set were 7 genes with *D. melanogaster* homologues that were functionally enriched for serine peptidase activity (up in ASG: *CG3916, CG32808, Ser6*; up in Starch: *lint, CG34458, γTry, Jon65Aiii*). Further to this enrichment, another *D. melanogaster* gene with a serine endopeptidase domain (CG6800) was DE, with higher expression in ASG abdomens. 4 transcripts with predicted annotations as *C. capitata* serine proteases were also called DE (up in ASG: GAMC01001732.1; up in Starch: XM_004517721.1, XM_004517825.1, XM_012306933.1). 5 *C. capita*ta transcripts with *D. melanogaster* homologues showed functional enrichment for the molting cycle (*Cht6, Lcp65Ab1* (*C. capitata* LCP5), *mey, neo, nyo*). Three of these (mey, neo, nyo) also showed enrichment for apical constriction. 3 *C. capitata* transcripts with *D. melanogaster* homologues contributed to the enrichment of the oxidative phosphorylation KEGG pathway (*SdhA, Vha100-2, Vha100-4*).

Beyond the transcripts described by functional enrichment, there was a wide range of additional *C. capitata* transcripts with *D. melanogaster* homologues showing DE. Of particular interest was a transcription factor associated with behaviour in *D. melanogaster* (*acj6*), which is associated with odorant receptor gene expression, as well as another transcription factor (CG15073). Given the dietary selection imposed, it was also interesting to observe DE in *ppk28*, a gene involved in response to water and osmoregulation in *D. melanogaster* (Cameron et al. 2010) and in Ent2, with reported roles in stress / stress signalling and associative learning in *D. melanogaster* (Knight et al. 2010). Predicted annotations of several of the *C. capitata* transcripts lacking *D. melanogaster* homologues was also interesting as within this set there were one odorant receptor (XM_004534136.1 (LOC101450382, Odorant receptor 67a-like)) and one odorant binding protein (XM_004521128.2 (Obp99a)) and the *C. capitata* gene CYP4E6 (AF028819.1), a cytochrome P450 associated with the processing of extrinsic and intrinsic metabolites (Danielson et al. 1999).

### VALIDATION BY QRT-PCR

Variation in gene expression as determined by mRNA-seq was validated using qRT-PCR (Table S1, S4). There was extremely close agreement between qRT-PCR and mRNA-seq expression. The pattern of gene expression was the same in all replicates of all 15 genes of interest (GOI) tested (Fig. S3-S17; Table S4) except in one case (GAMC01008957.1, AG1HT/SG1HT, Fig. S10). Statistical testing showed significant (*P* < 0.05) differences in 2^-ΔCT^ values between ASG and Starch flies in 14 out of 20 tests (Table S4). Within the set of validated GOIs, genes matching *D. melanogaster* homologues, such as SER1 (GAMC01003520.1) and TRY7 (GAMC01004081.1) were also validated. Overall, the qRT-PCR experiments provided a strong validation of the bioinformatics analysis of the mRNA-seq data, with 119 out of 120 qRT-PCR tests showing the same direction and/or magnitude as the mRNA-seq.

### MATE CHOICE TESTS

At generation 60, homotypic pairs predominated (Fig. 2, Table S5) and S males mated most frequently (Fig. 2, Table S5). However, at generation 90 these patterns were no longer evident. The results are described in more detail, below.

**Figure 2.**
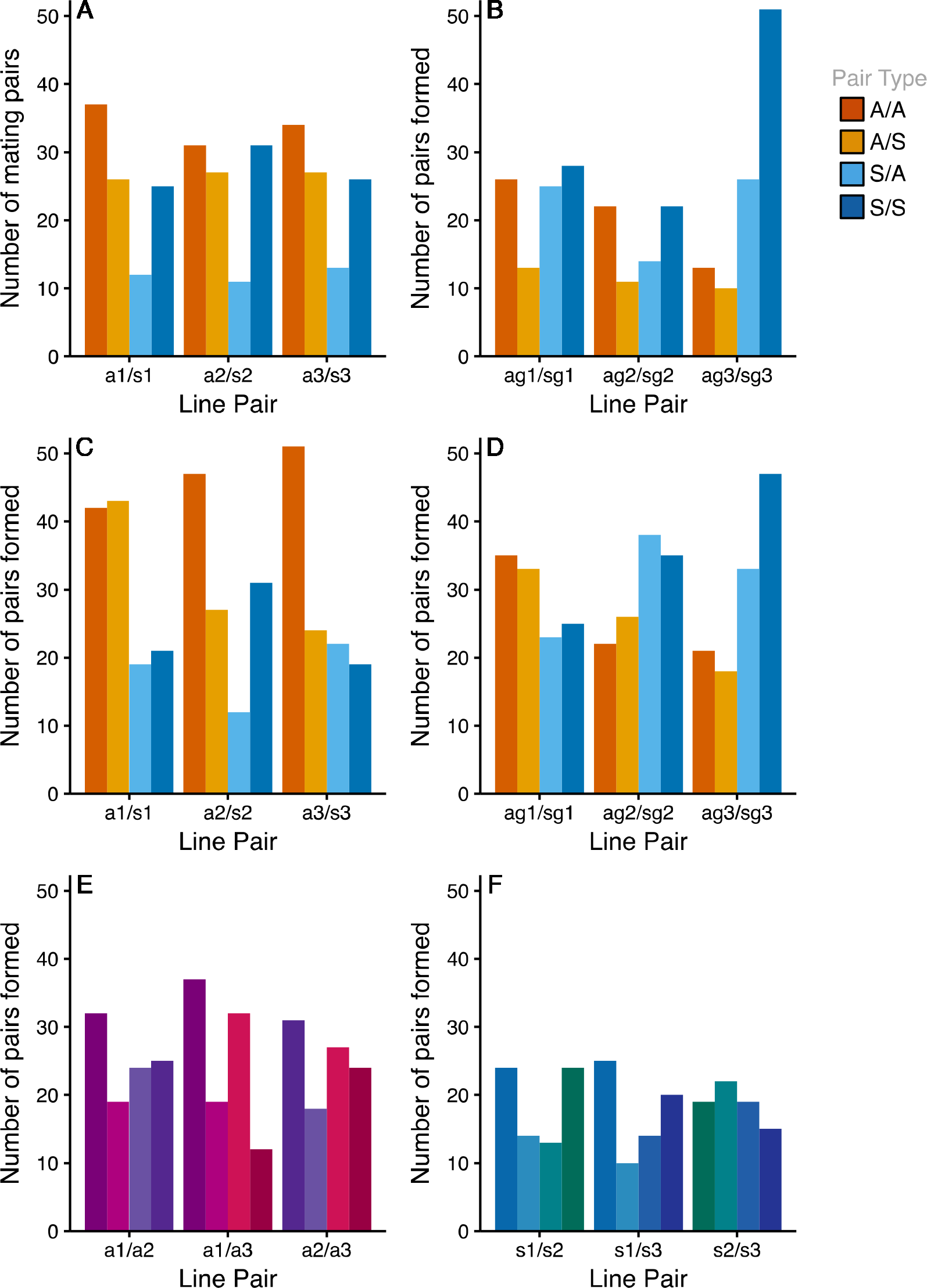
The number of mating pairs formed in multiple choice mating tests between ASG and Starch dietary selection lines after 60 & 90 generations of selection. Each plot shows three replicates, with lowercase letters representing diet (ASG = a or Starch = s) and replicate information (1-3). Upper case letters represent ‘pair types’ formed within that replicate (homotypic = AA, SS; heterotypic = AS, SA). Dark Orange bars represent homotypic pairings between the ASG males and females. Light orange bars represent heterotypic pairings composed of a male from ASG, and a female from Starch. Light blue bars represent the opposite heterotypic pairing, Starch male and ASG female. Dark blue bars represent homotypic matings between a Starch male and female. A) Flies reared on their own larval diet, tested at 60 generations. B) Flies reared on a common garden diet for two generations prior to testing, tested at 60 generations. C) Flies reared on their own larval diet, tested at 90 generations. D) Flies reared on a common garden diet for two generations prior to testing, tested at 90 generations. E) Flies from ASG line replicates tested against other ASG line replicates. Colours represent hetero- and homotypic matings by the same light / dark pattern as above, but with an individual colour pair for each line replicate. All flies reared on their own larval diet, tested at 90 generations. F) Flies from Starch line replicates tested against other Starch line replicates. Colouration as in E. All flies reared on their own larval diet, tested at 90 generations.

#### Sexual isolation (PSI), sexual selection (PSS) and total isolation (PTI) indices

At generation 60, the mating tests for individuals tested on their own diets showed significant deviations from random PTI (Table S6, S7) driven by significant deviations from random PSI (Table S6, S7). There was a clear pattern for homotypic (diet assortative) pairings to exhibit PSI > 1, and heterotypic (diet dissasortative) pairings to show negative values. The resulting PTI was significantly >1 for ‘AA’ mating type pairs in 2/3 replicates. Also, PTI coefficients for the ‘AB’ mating pair types were significantly < 1 for all lines (0.44 - 0.52). When tested in the common garden, the patterns were similar. In 2/3 replicates (Table S6, S8) the common garden environment removed any significant effect of treatment on PSI. PSS showed no significant differences from one in 2/3 replicates. PTI was significantly < 1 for ‘BA’ mating types in all line replicates (0.4 - 0.64). Hence at generation 60 there was evidence for significant assortative mating by diet in both the ‘on diet’ and common garden tests. A competitive advantage of ASG males was evident ‘on diet’, mediated by sexual selection. The maintenance of the assortative mating pattern in the common garden suggested that genetic adaptation within the lines according to their different environments resulted in the evolution of incipient divergence.

At generation 90, mating tests on individuals tested on their own diets again showed significant deviations from random PTI (Table S6, S9) driven by significant deviations from random PSI in a single replicate (Table S9). However, all line replicates showed significant deviations from random PSS (Table S9), indicating no recapture of the assortative mating by diet that was observed at generation 60. This lack of replication of the earlier pattern was also apparent when flies from generation 90 were tested after rearing on a common garden diet, with only a single line showing any significant deviation from random PTI, PSI or PSS (Table S6, S10), and as this effect was not driven by any deviation from random PSI, it was not likely associated with assortative mating by diet. At generation 90 the line replicates were also tested against each other within their dietary regimes, to test directly for potential confounding effects of genetic drift on any sexual isolation observed. Within the ASG regime, significant deviation from random PTI was seen in only one replicate (Table S6, S11) and again was driven by significant deviation in PSS and not PSI, suggesting a lack of assortative mating. In the Starch regime there was no significant deviation from PTI (Table S11).

#### Analysis of isolation - I_PSI_ - Isolation index

At generation 60, all lines showed significant assortative mating when tested on their own (*X*^2^ = 27.03, d.f. = 6, *P* < 0.001) and on the common garden (*X*^2^ = 19.7, d.f. = 6, *P* = 0.003) diets, (Table S12). However, at generation 90 this signature of assortative mating was not observed in own or common garden diets (Fishers Combined test, *X*^2^ = 5.3, d.f. = 6, *P* = 0.51). When lines within dietary regimes were tested, there was no significant signature of isolation for the ASG regime (*X*^2^ = 7.5, d.f. = 6, *P* = 0.26) but a significant isolation index in Starch (*X*^2^ = 19.3, d.f. = 6, *P* = 0.004).

#### Asymmetry in matings - IA_PSI_ - Asymmetry index

*IA*_PSI_ indicated any asymmetry in the proportion of homo- versus heterotypic matings in the own diet tests. In generation 60, *IA*_PSI_ was consistently significantly < 1 for heterotypic pairs (‘BA’ mating type), showing a consistent effect of diet across replicates (Fisher’s *X*^2^ = 23.22, d.f. = 6, *P* < 0.001). The frequency of heterotypic pairs was skewed towards BA type matings, i.e. those comprising S females and ASG males. No significant asymmetry between the proportions of homo-versus heterotypic matings was observed in generation 90.

#### Divergence in W - Cross product estimator of sexual selection

In the final analysis we calculated *W*, fixed at one for the fittest individual of each sex. There was no significant difference between treatments in female specific *W* at any generation 60. However, *W* for S males was lower than for ASG males in generation 60 (Fishers *X*^2^ = 24.28, d.f. = 6, *P* < 0.001). This pattern was also recovered in generation 90 (*X*^2^ = 15.75796, d.f. = 6, *P* = 0.015). However, ASG females also had significantly a significantly higher *W* than S females across all replicates.

## Discussion

We observed evolved differences in the expression patterns of genes involved in metabolism, development and OXPHOS in experimentally evolved populations of the medfly subjected to divergent nutritional selection. These differences underlie the phenotypic signatures of adaptation previously described (Leftwich et al. 2017). However, this adaptation was not associated with any consistent assortative mating by diet between the populations. mRNA-seq successfully captured variation in gene expression in sexually mature males from both selection diets. As all flies were reared on a common garden diet prior to RNA extraction, the measurement of evolved gene expression differences was maximised and proximate responses to diets minimised. The mRNA-seq results for each diet regime and body part were highly repeatable across independent biological replicates. qRT-PCR provided strong support for the genome-wide patterns of gene expression in the mRNA-seq. 214 transcripts showed DE above 2 log_2_OFC between treatments. Within this set, 94 transcripts were matched to homologous genes in *D. melanogaster* (44%). The functional description of these homologues, alongside the annotations available for the medfly showed divergence in gene expression patterns between sexually mature adult males from both nutritional regimes.

The larval diets which defined the ASG and Starch regimes were highly different in terms of both caloric value and specific nutritional content, giving insight into the likely selection pressures to which each population was challenged. The ASG larval diet contained over twice the Kcal/L in comparison to the Starch. Also, the addition of corn meal to the ASG diet offered additional sources of carbohydrates, proteins, and other dietary nutrients (http://ndb.nal.usda.gov/). This dietary complexity is important as it is the specific nutritional content of diets, rather than caloric content *per se*, that is known to affect life history traits such as lifespan (Mair et al. 2005). The effects of selection on these diets are evidenced by evolved changes in male body mass at sexual maturity (Leftwich et al. 2017).

The divergent nutritional selection resulted in a pattern of evolved differences in gene expression. Given the major differences between the diets in the complexity and diversity of carbohydrate content, DE in some form of nutrient metabolism was expected. Indeed, homologues of DE genes in the abdomen tissues of males were functionally enriched for ‘Serine-type endopeptidase activity’. The finding that different suites of these peptidases, (putative digestive enzymes; Ross et al. 2003) were upregulated in the different lines could suggest differentiation in digestive strategy as a basis for adaptation to diet. Several Drosophilid systems that exhibit divergence driven by host specialisation show similar, although more extensive, patterns of DE in nutrient related metabolic genes (e.g. Dworkin and Jones 2009; Wurmser et al. 2011; Etges 2014; Etges et al. 2015; Guillén et al. 2015). This effect is predicted on the basis that nutrients available in the larval diet can have large effects on survivorship and also have the potential to drive selection for optimal ability to utilise novel host nutrients. Within the medfly, the impact of larval rearing diet on male adult life history is substantial (e.g. Arita and Kaneshiro 1988; Kaspi et al. 2002). DE in metabolic genes could be associated with phenotypes manifested through effects on body size and nutrient reserves (Davidowitz and Nijhout 2004; Edgar 2006), but also through the potential for metabolic programming of adult metabolic traits by larval conditions (Matzkin et al. 2013; May et al. 2015).

Interestingly, divergent DE was observed in genes associated with OXPHOS in the Starch regime flies. In the abdominal tissue of flies from a Starch selective background, three genes with homologues enriching to the oxidative phosphorylation pathway (ko00190) were expressed at higher levels despite being reared on a common garden diet. Genes involved in OXPHOS are expressed in, or interact with the mitochondria which, due to their role as the centres of cellular energy production and rapidly evolving independent genome (mtDNA), are thought to be essential genes in adaptation and speciation (Gershoni et al. 2009; Ballard and Melvin 2010). An important example of this is the lake whitefish species complex (*Coregonus* spp.), in which divergence in OXPHOS gene expression in adult fish shows a tight relationship to the energetic phenotypes represented by two reproductively isolated morphs (Trudel et al. 2001; Derome et al. 2006; St-Cyr et al. 2008; Nolte et al. 2009; Jeukens et al. 2010; Renaut et al. 2010; Evans and Bernatchez 2012). The tissue specific DE seen in OXPHOS genes between males from ASG and Starch backgrounds may be associated with behavioural divergence (courtship activity) in these lines (Leftwich et al. 2017). Starch males showed elevated levels of courtship behaviours related to the expression of pheromones (Leftwich et al. 2017) and the up regulation of energy-related genes in the Starch male abdomen is consistent with this and could reflect the energetically costly process of pheromone biosynthesis (Dicke and Sabelis 1988; Jurenka 2004). The capture of signatures of changes to OXPHOS genes is interesting because it is believed to be important in responses to novel diets through detoxification and altered patterns of host related hormone regulation (Dworkin and Jones 2009; Wurmser et al. 2011).

Olfactory and gustatory receptor (OGR) genes are often associated with population divergence by host specialisation (McBride and Arguello 2007; Wurmser et al. 2011; Etges 2014) and the establishment of prezygotic barriers during speciation (Smadja and Butlin 2009). Such genes mediate chemosensory responses and are associated with host recognition, but also perception of pheromonal communication (Galindo and Smith 2001). Hence it was interesting to observe DE in an OGR gene in this study (OR67a-like), as well as two olfactory binding proteins (OBPs) (Obp19d, Obp99a). Patterns of DE in similar OBPs are seen between allopatric populations of *D. mojavensis* adapted to different host cacti and, in conjunction with other OBPs and behavioural genes, have been suggested to contribute to further population divergence (Etges 2014).

Given the previous evidence for nutritional adaptation (Leftwich et al. 2017) the finding that, at 60 generations, there was a consistent pattern of assortative mating by diet across the regimes was an interesting one. However, the tests at generation 90 did not replicate this pattern. In principle the two larval rearing diets provided sufficiently distinct selective environments to drive the evolution of parallel divergence (Leftwich et al. 2017). However, in this experiment, this did not lead to the establishment of a consistent signal of RI between populations. This may have been expected, as even geographically isolated natural populations have not returned significant RI across a global scale (Cayol et al. 2002; Lux et al. 2002). Despite this, significant behavioural differences have been observed between similarly geographically isolated populations, in courtship song (Briceño et al. 2002), and also courtship behaviour (Briceño et al. 2007; Diamantidis et al. 2008b). Further differentiation has been observed between global populations in other life history traits such as growth rate, longevity, and sexual maturation (Diamantidis et al. 2008a, 2009), as well as pre-adult traits (Diamantidis et al. 2011b), and resilience to domestication (a potential proxy for an enforced change of host) (Diamantidis et al. 2011a).

The variance seen across these traits in global populations of medfly reflects overall genetic diversity of these populations, which is closely linked to the medfly’s invasion history (Gasperi et al. 2002; Karsten et al. 2015). Lack of divergence (isolation) between populations that vary globally is likely due to the effects of gene flow between populations (Malacrida et al. 1998, 2007; Karsten et al. 2015), potentially mediated by invasions due to human transport (Wilson et al. 2009). This migration between populations has likely suppressed the capacity for specialisation that can exist, as shown here, and has instead maintained the plasticity exhibited by the medfly as a generalist, in both host and temporal space (Yuval and Hendrichs 2000). Despite this, this study demonstrates that a laboratory population (over 20 years isolated from the wild) maintained sufficient genetic variation to respond to experimentally-imposed divergent selection. Relating this to natural populations, as quarantine measures become increasingly effective and reduce gene flow between global medfly populations (Karsten et al. 2015), the adaptive potential observed here may lead global populations to diverge further.

## ACKNOWLEDGEMENTS

We thank Damian Smith and Emily Fowler for valuable guidance and problem solving in the laboratory tests and qRT-PCR, Phil Leftwich and Lucy Friend for inventive discussions on experimental design. We thank the BBSRC (BB/L003139/1 grant to TC and IM) and the NERC (iCASE PhD studentship to TC and WN) for funding.

## DATA ARCHIVING

Data are available on the GEO database (Barrett et al. 2013): GSE86029; GSM2291180 to GSM2291191.

## AUTHOR CONTRIBUTIONS

W.N., I.M. and T.C. conceived the research; W. N. conducted the experimental work; W.N and I.M. analysed the data; W.N., I.M. and T.C. wrote the paper.

## Supporting Figures and Tables

**Figure S1.** Distribution of expression levels before and after normalisation. Left panel represents the raw non-redundant reads, right panel represents the expression levels generated following the application of subsampling normalisation followed by quantile correction. On the X axis are the samples and, on the Y, the log2 expression levels with an offset of +20.

**Figure S2.** Hierarchical differential expression (DE) histogram. The frequency distribution of log2OFC DE between sum expression in tissue types (HT/Ab), and in treatment (A/S). The distribution of DE between treatment types, in red, falls below the distribution of DE between tissue types, in black.

**Figures S3 - S7.** Validation on RNA-seq analysis using RT-qPCR for genes expresses in both HT and Ab Tissue. Each plot consists of a panel of six figures. The left- and right-hand columns are expression profiles of the candidate gene of interest (GOI). These expression profiles show the algebraic sum of abundances of incident reads at every position of the reference transcript, for each treatment. Expression levels from ASG tissue is presented in the left-hand column, expression levels from the Starch tissue in the right-hand column. Head /Thorax (HT) tissue is presented in the top row of both columns, and in blue. Abdominal (Ab) is presented in the bottom row of both columns, and in red. Line replicates are indicated by different line types. The central column of two figures shows the normalised expression level following analysis (top), and the 2^-ΔCT^ expression level suggested by low throughput validation using RT-qPCR. Again, ASG left, Starch right, HT blue, Ab red.

**Figures S8 - S12.** Validation on RNA-seq analysis using RT-qPCR for genes expresses in HT tissue only. Layout of these plots is as described for Fig. S3 – S7.

**Figures S13 - S17.** Validation on RNA-seq analysis using RT-qPCR for genes expresses in Ab tissue only. Layout of these plots is as described for Fig. S3 – S7.

**Table S1. Primers used for RT-qPCR Validation.** Forward (f) and reverse (r) primers used for each GOI in the RT-qPCR validation experiment.

**Table S2. Quality control and genome matching of RNA-seq data used in this study.** ‘fastq2fasta’ describes the conversion of sequence data FASTQ format to FASTA format, ‘%Acc’ describes the accuracy of this conversion. ‘R2NR’ denotes the conversion of the FASTA files into non-redundant format (see text), ‘NR’ = non-redundant, ‘R’ = redundant, ‘C’ = complexity (see text). ‘Bstrp’ describes the 29M fixed total subsampling of the data, conducted as a normalisation step (see text).

**Table S3a. Differentially expressed (DE) transcripts (full dataset) found in Head/Thorax tissue** taken from sexually mature males after 60 generations of experimental evolution on one of two dietary treatments: ASG (AG) or Starch (SG)(see text). Flies were reared for two generations on a common garden diet prior to flash freezing at 7 days post eclosion and subsequent RNA extraction. Transcripts are shown with their NCBI identifier. Normalised expression values are shown for each replicate of each treatment, these are labelled treatment/replicate/sex/tissue (e.g. AG / 1 / M / Ab). DE was called as log2 of the difference between either the highest or lowest (+ 20) normalised expression value of the three treatment replicates, depending on which treatment showed higher expression overall. Also shown are the *Drosophila melanogaster* homologues identified for each gene using tBLASTx (see text). Further to this NCBI annotation information is given.

**Table 3b. Differentially expressed (DE) transcripts (dataset with duplicate entries removed) found in Head/Thorax tissue** taken from sexually mature males after 60 generations of experimental evolution on one of two dietary treatments: ASG (AG) or Starch (SG)(see text). Dataset as described in Table S2a, but where multiple transcripts were present in the NCBI set used to map RNA-seq read to, all reference sequences were aligned to ensure similarity, and duplicates were removed, in these cases the transcript which exhibited the highest DE value is presented as exemplar, and the NCBI id of duplicates is also given.

**Table S3c. Differentially expressed (DE) transcripts (full dataset) found in Abdominal tissue** taken from sexually mature males after 60 generations of experimental evolution on one of two dietary treatments: ASG (AG) or Starch (SG)(see text). Flies were reared for two generations on a common garden diet prior to flash freezing at 7 days post eclosion and subsequent RNA extraction. Transcripts are shown with their NCBI identifier. Normalised expression values are shown for each replicate of each treatment, these are labelled treatment/replicate/sex/tissue (e.g. AG / 1 / M / Ab). DE was called as log2 of the difference between either the highest or lowest (+ 20) normalised expression value of the three treatment replicates, depending on which treatment showed higher expression overall. Also shown are the *Drosophila melanogaster* homologues identified for each gene using tBLASTx (see text). Further to this NCBI annotation information is given.

**Table S3d. Differentially expressed (DE) transcripts (dataset with duplicate entries removed) found in Abdominal tissue** taken from sexually mature males after 60 generations of experimental evolution on one of two dietary treatments: ASG (AG) or Starch (SG)(see text). Dataset as described in Table S2c, but where multiple transcripts were present in the NCBI set used to map RNA-seq read to, all reference sequences were aligned to ensure similarity, and duplicates were removed, in these cases the transcript which exhibited the highest DE value is presented as exemplar, and the NCBI id of duplicates is also given.

**Table S4. Statistical validation of RT-qPCR results.** Results of Welch’s two sample t test conducted on the 2-ΔCT values of qRT-PCR validation experiments for 15 GOI. Genes were selected to test both tissue specific expression patterns, or organism wide expression. Genes which did not return signal (below detection limit, highlighted in pink) were assigned a CT value of 40 to facilitate statistical testing.

**Table S5. Sample size (n) and total recorded matings, also broken down by mating pair type (Female/Male: AA, AB, BA, BA), at generations 60 & 90.** Raw frequencies shown. Treatments represent: ASG/ASG - Individuals from the ASG regime, tested on the ASG larval diet, ASG/S - Individuals from the ASG regime, tested after two generations of rearing on the Starch larval diet, S/S - Individuals from the Starch regime, tested on the Starch larval diet, S/ASG - Individuals from the Starch regime, tested after two generations of rearing on the ASG larval diet, ASG/G - Individuals from the ASG regime, tested after two generations of rearing on the common garden (Glucose) larval diet, S/G - Individuals from the Starch regime, tested after two generations of rearing on the common garden (Glucose) larval diet.

**Table S6. G statistics for generation 60 and generation 90 mating tests.** G statistics calculated within JMATING (see text) to test for deviations from random mating across the whole coefficient dataset. GI = G test of PSI coefficient, GS = G test of PSS coefficient, GT = G test of PTI coefficient.

**Table S7. Multiple choice mating tests with all flies reared on their own larval diet, tested at generation 60.** Pair sexual isolation coefficient (PSI), pair sexual selection coefficient (PSS), and pair total isolation coefficient (PTI), calculated after Rolan-Alvarez and Caballero (2000), for each line pairing, and possible mating pair type. Coefficient values represent mean coefficient values generated after 10,000 bootstrap resamples from the observed (for PTI) or observed and expected frequencies (for PSI and PSS). Stars represent a significant bootstrap probability of rejecting the null hypothesis, that the coefficient is not different from 1 (no preference, or random mating). Lowercase letters represent diet (ASG = a or Starch = s) and replicate information (1-3). Upper case letters represent ‘pair types’ (homotypic = AA, BB; heterotypic = AB, BA).

**Table S8. Multiple choice mating tests with all flies reared on a common garden glucose based larval diet, tested at generation 60.** Pair sexual isolation coefficient (PSI), pair sexual selection coefficient (PSS), and pair total isolation coefficient (PTI), calculated after Rolan-Alvarez and Caballero (2000), for each line pairing, and possible mating pair type. Coefficient values represent mean coefficient values generated after 10,000 bootstrap resamples from the observed (for PTI) or observed and expected frequencies (for PSI and PSS). Stars represent a significant bootstrap probability of rejecting the null hypothesis, that the coefficient is not different from 1 (no preference, or random mating). Lowercase letters represent diet (ASG = a, Starch = s, Glucose = g) and replicate information (1-3). Upper case letters represent ‘pair types’ (homotypic = AA, BB; heterotypic = AB, BA).

**Table S9. Multiple choice mating tests with all flies reared on their own larval diet, tested at generation 90.** Pair sexual isolation coefficient (PSI), pair sexual selection coefficient (PSS), and pair total isolation coefficient (PTI), calculated after Rolan-Alvarez and Caballero (2000), for each line pairing, and possible mating pair type. Coefficient values represent mean coefficient values generated after 10,000 bootstrap resamples from the observed (for PTI) or observed and expected frequencies (for PSI and PSS). Stars represent a significant bootstrap probability of rejecting the null hypothesis, that the coefficient is not different from 1 (no preference, or random mating). Lowercase letters represent diet (ASG = a or Starch = s) and replicate information (1-3). Upper case letters represent ‘pair types’ (homotypic = AA, BB; heterotypic = AB, BA).

**Table S10. Multiple choice mating tests with all flies reared on a common garden glucose based larval diet, tested at generation 90.** Pair sexual isolation coefficient (PSI), pair sexual selection coefficient (PSS), and pair total isolation coefficient (PTI), calculated after Rolan-Alvarez and Caballero (2000), for each line pairing, and possible mating pair type. Coefficient values represent mean coefficient values generated after 10,000 bootstrap resamples from the observed (for PTI) or observed and expected frequencies (for PSI and PSS). Stars represent a significant bootstrap probability of rejecting the null hypothesis, that the coefficient is not different from 1 (no preference, or random mating). Lowercase letters represent diet (ASG = a, Starch = s, Glucose = g) and replicate information (1-3). Upper case letters represent ‘pair types’ (homotypic = AA, BB; heterotypic = AB, BA).

**Table S11. Multiple choice mating tests with all flies reared on a common garden glucose based larval diet, tested at generation 90 - WITHIN TREATMENT.** Pair sexual isolation coefficient (PSI), pair sexual selection coefficient (PSS), and pair total isolation coefficient (PTI), calculated after Rolan-Alvarez and Caballero (2000), for each line pairing, and possible mating pair type. Coefficient values represent mean coefficient values generated after 10,000 bootstrap resamples from the observed (for PTI) or observed and expected frequencies (for PSI and PSS). Stars represent a significant bootstrap probability of rejecting the null hypothesis, that the coefficient is not different from 1 (no preference, or random mating). Lowercase letters represent diet (ASG = a or Starch = s) and replicate information (1-3). Upper case letters represent ‘pair types’ (homotypic = AA, BB; heterotypic = AB, BA).

**Table S12. *I*_PSI_ isolation index values (Coyne et al. 2005) for each line pairing after 60 generations of selection.** Values presented are bootstrap means with standard deviations; P values are the two-tail bootstrap probability that the value is significantly different from one (equivalent to random mating), after 10,000 iterations of resampling.

